# “Sulcus Sink”: A Compact Binary and Semi-Automated Inverse Dijkstra-based System for Describing Sulcal Trajectories

**DOI:** 10.1101/2020.02.18.955096

**Authors:** Rudolph Pienaar, Christian Hasselgrove, Kiho Im, David Kennedy, P Ellen Grant, Denise Boriel, Lena Tang, Nikos Makris

## Abstract

We present a description of a system that uses a compact binary representation to describe and trace sulci on a reconstructed human cortical surface, based on a set of human-generated targets. The inputs to the system were manually created on a training set of 20 normal subjects (11 females, 9 males) with ages 22 – 40 years. T1 weighted MPRAGE images were collected on a Siemens 3T Trio scanner, with TR/TE = 2530/3.3, matrix = 256×256, FOV = 256mm, slice thickness 1.33mm. The resultant images were reconstructed with Freesurfer, and 10 sulci on each hemisphere were traced by an expert human operator and independently assessed for accuracy. Presented with these input trajectories in its training phase, the system attempted to determine a compact binary feature vector of each sulcus on each subject using as descriptor a binary parametrized function of several surface-geometry variables (such as mean curvature, sulcal depth, edge length, etc.). This function was optimized in a supervised learning fashion using a Dijkstra-based graph theory formulation, in which the binary weights were used to define graph edge costs. In the setup phase, the system was presented with sulcal trajectories already defined on surfaces, and then adjusted its parametrized weights in a binary fashion to minimize differences between the training input path and its Dijkstra-generated output path. Once the setup phase was complete and sulci had been described in a per-sulcus, per-subject manner, we generalized the per-sulcus description across all the subjects to construct a template binary word for each specific sulcus. The performance of the system for each subject and each sulcus, and for each template sulcus group was measured against the original human reference in both a quantitative and qualitative manner. Individual subjects generally showed very good optimization to their manually traced training samples across all sulci, with 91% average overlap within 4mm of the human target. Generalized group results, as expected, showed less overlap with the original human targets, but still performed on average with 80% overlap. Quantitatively, the group results were nonetheless for the most part quite acceptable to an independent human evaluator. The parametrized binary weight description that drives the Dijkstra path optimization is presented as a mechanism to succinctly and compactly describe individual human sulci and groups of sulci.

## I. Introduction

This paper presents a description of a methodology called “sulcus sink” that has been developed at the Massachusetts General Hospital’s (MGH) Center for Morphometric Analysis (CMA). Sulcus sink attempts to describe sulci using a compact binary parametrized description. Presented with a human generated reference sulcal trace, the method searches in a binary fashion through a parametrized weight space of graph edge costs and uses Dijkstra’s algorithm to optimize the sulcal traces that result from a given weighting. The system attempts to find a weight parametrization that minimizes the error between the human reference and Dijkstra-generated trajectories.

This methodology arose from the very practical need to aid human operators as they manually traced sulcal lines on reconstructed surfaces. The operational use of the system would allow human operators to simply and relatively quickly select sulcal start and terminal points on a reconstructed surface. The sulcus sink system, trained on a cohort of human generated references, would complete the sulcal trace, based on a generalization of earlier optimization analyses.

The purpose of the system is therefore not to automatically extract sulci, but rather to attempt to capture the relative weighting that human operators might place on components of cortical geometry as they generate manual traces, and encode this information in a compact binary word. An important implication of such a weight parametrization description is that it allows us to attempt to propose unbiased sulcal-specific descriptions that have been generalized from an underlying set of per-subject and per-sulcus optimal parametrization. Given such a binary parametrization that weighs graph edges between vertices of the reconstructed cortical mesh, the optimization engine searching through this graph-based representation naturally lends itself to processing by Dijkstra’s algorithm [7]. The system is considered “semi-automated” in that it does require, for a given sulcus, start and terminus points. A complete description, therefore, is the binary parametrization, sulcus start and terminus points, and Dijkstra’s algorithm on resultant edge costs.

While the emphasis of the paper is on the technical and operational aspects of the methodology itself, preliminary results on the semi-automated tracing of sets of twenty sulci on twenty subjects is also presented. Based on the characteristics of individually optimized sulcal traces, we also attempt to generalize over all similar sulci in all the subjects.

The automatic (or semi-automatic) tracing of sulci on the cortical surface is useful in a number of neurological applications, from surgical planning to cortical parcellation and morphometric analysis [10, 25, 16, 26, 9, 20], brain registration [13] and exploring brain development during pathologies [39].

A large volume of literature seeks to address the problem of automatically detecting sulci based around geodesics generated by various methods – from dynamic programming [17] using local shape measures (but still requiring the manual selection of sulcal start and end points), through the optimal calculation of geodesics on triangulated mesh surfaces [2] to generate isotropic geodesics. Such geodesics have also been computed by a skeleton extraction approach that solves an Eikonal equation [35] – itself a special type of a Hamilton-Jacobi equation [36]. A refinement to such isotropic geodesics has recently been extended to anisotropic conditions which are more highly adaptive to the anisotropic nature of cortical surface shape and which can lead to sulcal lines lying more accurately within sulcal valleys [34]. Statistical methods based on mean curvature values have also been considered [38] – these require surface registration and statistical training sets.

Geodesic approaches are intuitively appealing since they map to our understanding of the geometry of surfaces. Indeed, a geodesic is by definition a locally length minimizing curve. However, in order to better explore a more abstract parametrized description (including not only curvature, sulcal depth, and distance but also products of the same), we believe a more graph-theoretical approach wherein edge weights can be arbitrarily (and possibly less intuitively obviously) defined affords us better representational power. Such a graph-theoretical approach is ideally suited to Dijkstra’s algorithm.

The geodesic and statistical approaches for the most part attempt to solve or model geometric properties using a relatively small subset of available geometric properties – often the mean curvature. Extending these approaches to less intuitive geometric properties or considering the behavior of these systems under such circumstances is often not explored. Our approach here is to explicitly formulate our problem from the perspective of generating a compact binary weight description of multiple geometric properties, and to use this formulation as a descriptor of a given sulcus.

Using Dijkstra’s algorithm [7] in this manner as an optimizer for tracking human-traced sulcal paths is a relatively novel approach. In the context of cortical parcellation, Dijkstra’s algorithm has previously been used to calculate lengths on the cortical surface [3], and also as a basis for exploring geometric invariants in Hurdal’s analysis in classifying cortical sulci [22]. Conceptually, there are some broad similarities between our approach and Hurdal’s, however Hurdal attempts to describe sulci using invariant shape descriptors such as the writhe number, ropelength, and moment-based measures of a sulcal trajectory curve. Our work here parametrizes sulci not as complete curves, but rather as a vector of weights that define a cost function to a Dijskstra-based path search. The weight vector uses features calculated from the reconstructed cortical surface, primarily the mean curvature (using methods of discrete geometry [23][37][8]), distance between vertices, direction to end-point, etc.).

In some respects, this work touches on issues relating to the complexity of folded cortical surfaces. Several studies have previously attempted to quantify folding and/or complexity. Fractal dimension analysis has often been used as a technique for measuring “complexity” (14; 18; 15), as well as folding measures derived from cortical thickness analysis (41; 40) and metric distortions that arise from registering surfaces to an average template [42]. Although fractal measures are interesting in the abstract, we find them difficult tools to use as a means of understanding the topology of cortical folding, and hence sulcal paths. Numerous studies based on cortical thickness employ regional shape measures based on functions of mean curvature.

Other means of describing sulci using Mangin’s “sulcal roots” have also been proposed. These “sulcal roots” correspond to the first folding locations during antenatal life (4; 5; 27; 28; 29; 33; 1) and are objects derived from mean curvature minima and saddle points. The highly variable pattern of folding noted across adult brains can be recovered from successive scale-space analysis. Folding patterns are decomposed and their core embedded patterns simplified. Conceptually, Mangin’s work constructs “tree-like” structures tracing the development of simple folds to complex pattern at different scales. Our work is a complement to this approach. Sulcal roots provide a means for logically constructing how simple folds become complex patterns without focusing on the detailed curvature characteristics of the final patterns. Our work is less concerned with the exact location of the maximal sulcal depth and its branching pattern, and more focused on the surface topology which we analyze using several curvature functions.

Additional studies have attempted to consider the intrinsic geometry of the cortex from a more purely mathematical and computational basis [12], but followup studies in a similar vein using contemporary computing power have not been pursued. Moreover, these studies have focused on adult, not pediatric or developing newborn brain surfaces.

More recently, spherical wavelets have also used to quantify cortical folding [43, 31]. Development of surface folding was modeled through increasing wavelet powers and these wavelet coefficients were fitted to the Gompertz function, a model of self-limited growth. This allows predictions as the when sulcal trajectories might occur during development, and also provides measures as to the differential folding rate across developing surfaces. The Gompertz function has been used successfully in the past to describe brain volumetric growth (30; 19). A more direct methodology based on surface curvature features and functions of principle curvature has also been proposed to quantify the shape structure of the cortical manifold [32].

From an organizational perspective, Section II focuses on the experimental methods, describing the data set itself, details on the segmentation and the reconstruction process, and the sulci that were traced. Section III describes the Dijkstra based optimization process, and the sulcus sink system itself. The weight vector space, and search methods through this space are presented, and the creation of general templates from a set of optimal weights is considered. Section IV presents and discusses the results of the optimization, with specific examples of sulcal correlation to the training templates, as well as the generalization of the sulcal weights across all subjects. Finally, the paper is concluded in Section V.

## II. Experimental Methods

The Experimental Methods section presents and discusses the main data collection and preparation steps required by the sulcus sink system. Information specific to the data collection and preparation stream are presented, and the software tools that process the data are discussed.

### A. Data Collection and Surface Reconstruction

For this paper, twenty subjects were imaged. This population was age and gender balanced, with 11 males and 9 females, and ages ranging from 22 – 40 years old. All images were acquired on a 3T Siemens Trio system, T1 weighted MPRAGE images with TR/TE = 2530/3.3, matrix = 256×256, FOV = 256mm, slice thickness 1.33mm, and in-plane resolution of 1×1 mm^2^.

Once collected, these images were processed using Freesurfer [11, 6] and surfaces reconstructed. The gray/white junction (or simply, “white matter”) surface was used as the primary analysis surface. For each subject, each sulcus of interest was manually traced on an inflated representation of the white matter surface. FreeSurfer surfaces are tessellated wire-frame structures, typically consisting of more than 120,000 vertices for an average adult brain and provides an isotropic resolution of 1 mm^3^. In this study, the average vertex separation distance between individual mesh vertices was about 0.79mm. The mesh structure contains vertex-specific data, including: list of neighboring vertex indices; distance (in mm) to each neighboring vertex; Cartesian coordinates of each vertex in an anatomical space; and surface curvature along each link connecting a vertex to its neighbors.

Table I enumerates the sulci that were considered on each hemisphere, and Figure II.1 shows some manual traces on an example subject.

**Table I.**
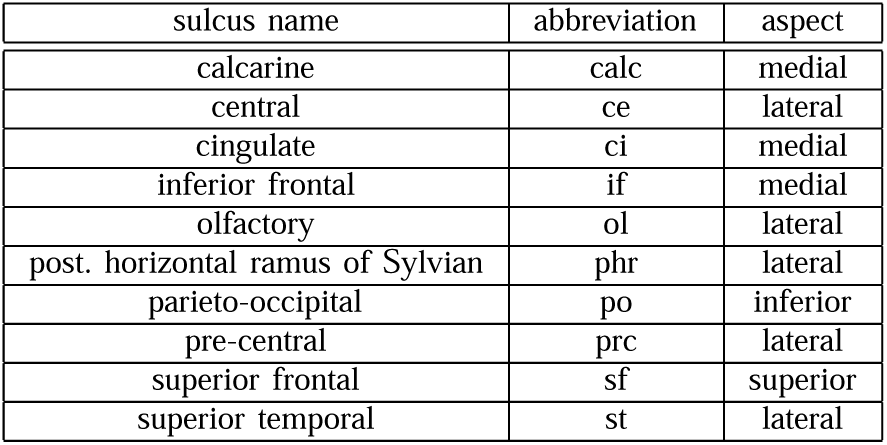
A List of sulci that were manually traced, and subsequently cost-function optimized.

| sulcus name | abbreviation | aspect |
| --- | --- | --- |
| calcarine | calc | medial |
| central | ce | lateral |
| cingulate | ci | medial |
| inferior frontal | if | medial |
| olfactory | ol | lateral |
| post. horizontal ramus of Sylvian | phr | lateral |
| parieto-occipital | po | inferior |
| pre-central | prc | lateral |
| superior frontal | sf | superior |
| superior temporal | st | lateral |

**Figure II.1.**
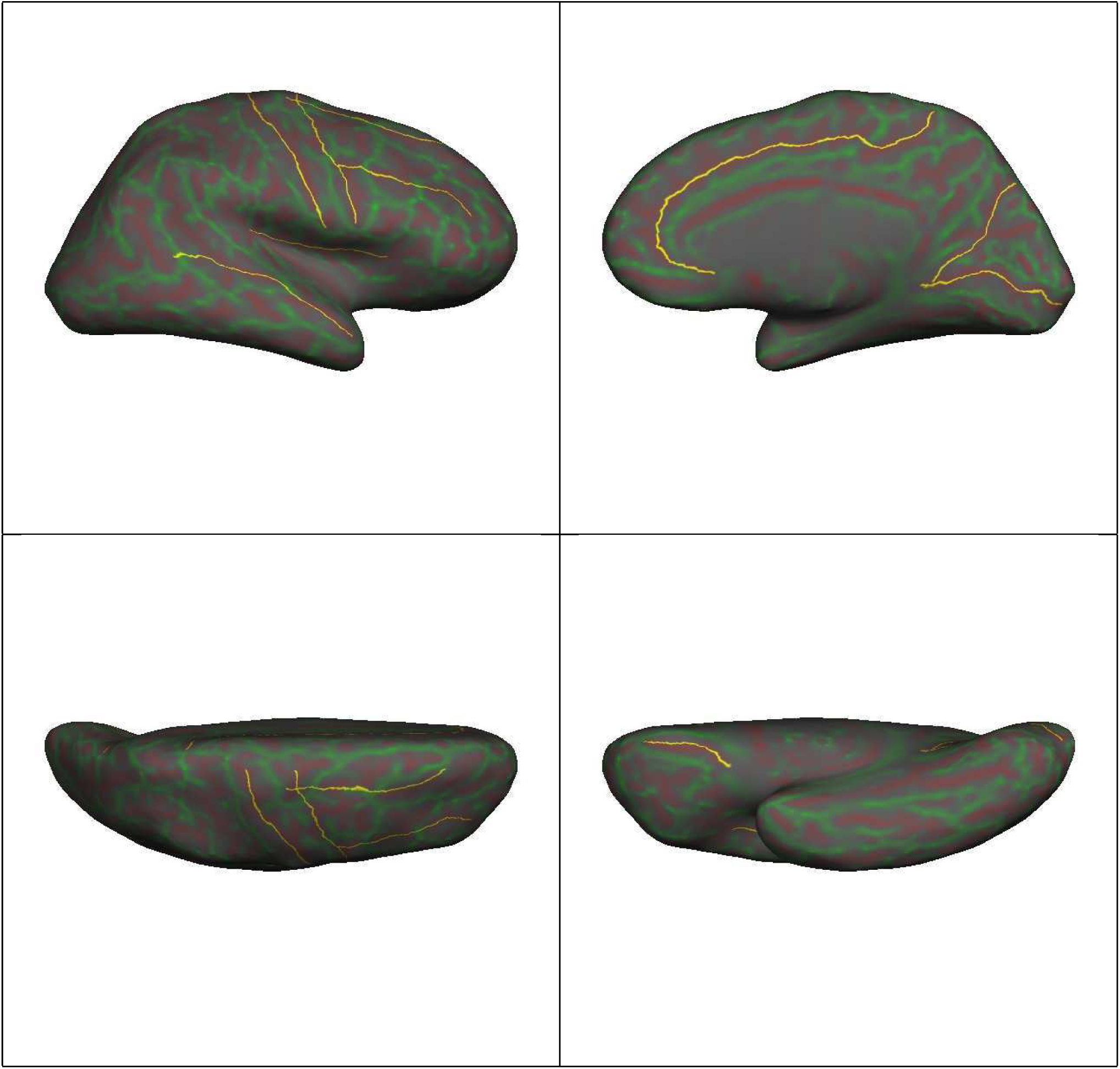
Sulci as traced by a human operator on a sample reconstructed right hemisphere inflated surface. In the top row, on left, is the lateral view. The sulci shown, from bottom and moving up in a clockwise direction: *sf, ce, prc, sf, if, phr*. On the top right is the medial aspect, dominated by the long *ci*, with the *po* (above) and *calc* (below) at the occipital pole. On the bottom left is the superior aspect showing the *ce* and *prc* (right to left) and the *sf* and *if* (top to bottom). The bottom right shows the *ol* on left. The same sulci were also traced on the contra lateral hemisphere. The background coloration of red and green denote curvature values of the white matter surface projected onto the inflated reconstruction. Red curvatures are positive denoting “inward” depression into the surface; green curvatures are negative, denoting “outward” bulges from the surface.

Initially, the system was “trained” in a supervised learning fashion that guided the optimization process. For each sulcus on each subject under consideration, an expert human operator manually traced the path of the sulcus using projections on the reconstructed brain surface. These paths were cross-checked against planar slices in a volumetric space to assure that they lay along anatomically correct parts of the sulcus [24].

## III. Optimization Methods

This section describes the actual optimization method used to minimize error between the human specified sulcal path, and the path generated by the “sulcal sink” software. It will describe the binary weight space, the cost function, and its overlap with the human reference from a Dijkstra graph-theory perspective.

### A. Dijkstra development and weight vector

The reconstructed surface structure generated by Freesurfer is a tessellated mesh that lends itself both visually and conceptually to a graph-theory based approach and in turn to processing by Dijkstra’s algorithm [7]. Dijkstra’s algorithm is a greedy algorithm that solves the single-source shortest path problem for a directed graph with non-negative edge weights. Essential components are (1) the fact that it is a “locally greedy” algorithm, i.e. operates under the assumption that a global optimum can be found by at each step in an iterative algorithm by choosing the local optimum, and (2) that connections (or weights) between nodes in the graph are non-negative values of some underlying cost function. Dijkstra’s algorithm will optimize for the lowest cost path in this connected graph.

Figure III.1 shows a part of the wire-frame mesh for a typical Freesurfer surface reconstruction, and a path running along this mesh. Given an *a priori* path connecting two points in a directed graph as shown, we attempt to find an edge weighting that maximizes the overlap between the given path and a Dijkstra optimization,

**Figure III.1.**
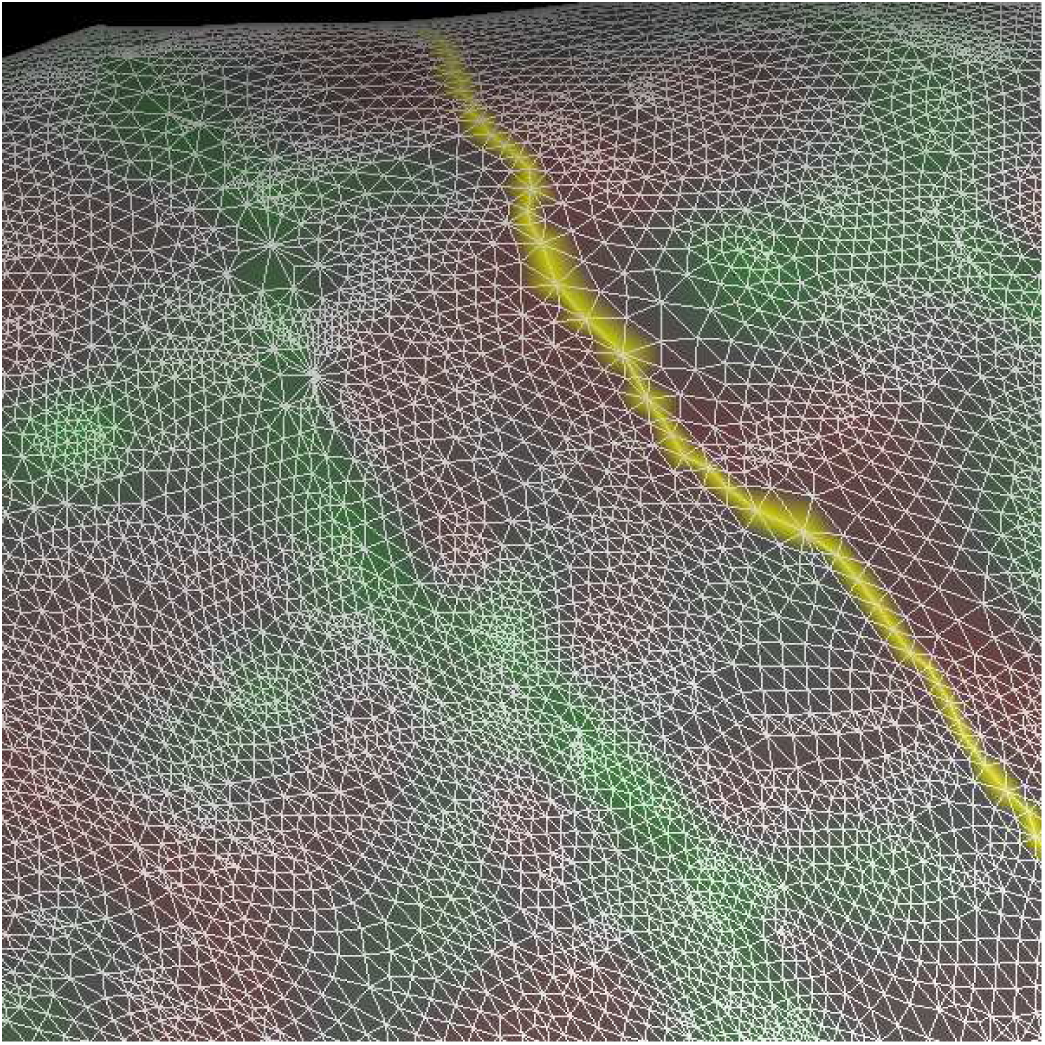
A path connecting two vertices on a Freesurfer surface mesh. The green and red colored faces denote “outward” and “inward” mean curvature respectively.

In the general case, we can create a cost function that is a multi-variable function that contains the vertex edge, a weight vector, and a conditional penalty matrix:

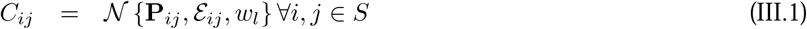

where P_*ij*_ is a penalty weight matrix, ℰ_*ij*_ the edge itself between each vertex point 𝒱_*i*_ and its neighbor 𝒱_*j*_, *w*_*l*_ a weight vector, and surface *S*. The cost function should have low (positive) values for graph edges that are deep within the sulcus, and higher cost values elsewhere. The connecting edge structure ℰ_*ij*_ contains several important fields, viz:

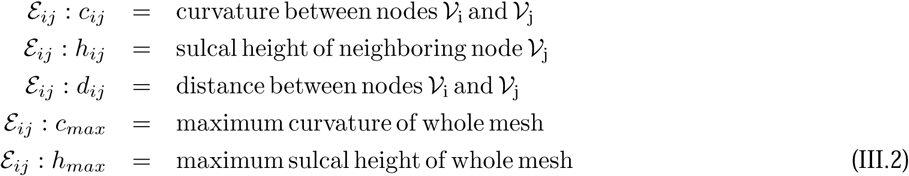

Within Freesurfer, both curvature and sulcal height are signed quantities. Since Dijkstra’s algorithm depends on non-negative cost values between nodes, we re-cast Freesurfer’s signed curvature and sulcal height values to be meaningful in the optimization context we are attempting to solve. This simply entailed shifting the sulcal displacement and surface curvature values upwards by the their respective maximum values for the whole surface:

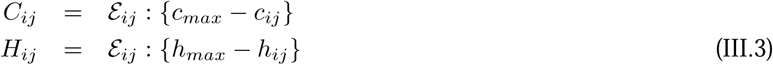

which casts high positive curvatures and sulcal heights close to zero and assigns highly negative curvatures and sulcal heights high positive values.

#### 1) Fundamental cost function parameters

For the purposes of this paper, we use three fundamental edge parameters, the edge length, *d*_*ij*_, the edge curvature *c*_*ij*_, and the edge sulcal height, *h*_*ij*_. Within a typical Freesurfer reconstruction, these parameters are all of the same magnitude – hence normalization of these values to some arbitrary reference was not necessary. Derived parameters are all the multiplicative combinations of these three fundamentals.

#### 2) Locally Greedy Component

Ignoring for the moment the non-locally greedy penalty matrix of the cost function in Equation III.1, we can express the locally greedy component as a product between the weight vector *w*_*l*_ and a set of edge parameters:

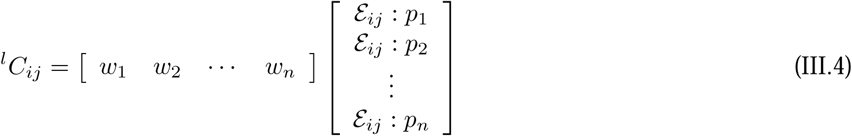

These edge parameters, dropping for notational simplicity the ℰ_*ij*_ : suffix, are described by the seven element set of all combinations of curvature, sulcal height, and distance, as well as an additional parameter, *p*_*ie*_, which is the alignment of the current edge direction *i* with the vector linking the current vertex node to the end vertex node *e*. The edge parameters are therefore (with reference to Equation III.2):

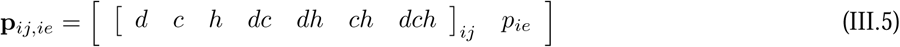

where compound parameters imply the product of each underlying parameter, i.e. *dc* implies distance times curvature. Note that the *p*_*ie*_ parameter is in fact 1 − *P*_*ie*_ where *P*_*ie*_ is the actual projection of the current edge the linking vector (for complete co-linear projection, *P*_*ie*_ = 1, which we would weigh as zero in a Dijkstra context, while a tangential direction would have zero projection, and hence a high *p*_*ie*_). Our locally greedy cost function is therefore:

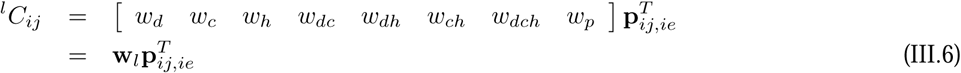

with the *ij* subscript denoting the parameters for the edge connecting vertex *i* and *j*, and the *ie* subscript denoting the direction from the current vertex *i* to the end vertex *e*. The first component of Equation III.6, the locally greedy edge weight vector w_*l*_, is the component that we seek to optimize.

#### 3) Penalty Matrix

The non-locally greedy component of the cost function is a penalty matrix is triggered by curvature transitions along the trajectory path. These arise when strict adherence to the underlying Dijkstra “greedy” behavior is in fact a bad strategy. This might arise if “bumps” or interruptions are found along a sulcal path - particularly when a gyrus “interrupts” or crosses over a sulcus. A purely locally “greedy” algorithm might route *around* this interruption instead of continuing *over* it to reach the remainder of the current sulcus. The non-linear component consists of a penalty matrix that toggles between unity and a penalty value based on any changes in curvature sign that are encountered along a path,

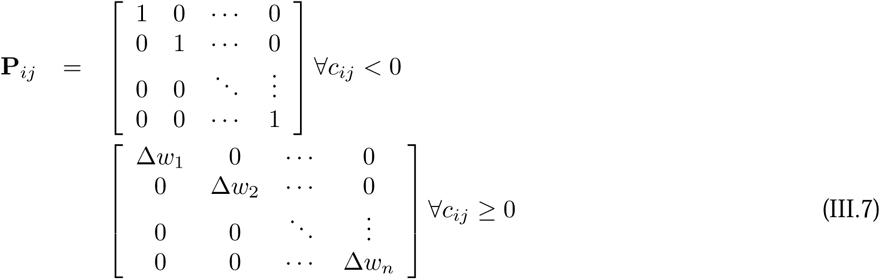

where the Δ*w*_*k*_ penalties weigh the original weight vector with an additional per-edge weight to counter the locally greedy nature of Dijkstra’s algorithm. The final cost function follows from Equations III.1, III.6, and III.7:

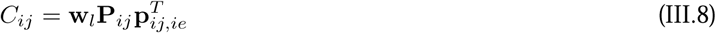

and has as diagonal penalty vector

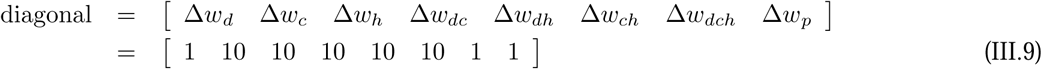

Since the penalty diagonal matrix was kept constant throughout, and only the weight vector was variable across a whole Dijkstra path, we can simplify Equation III.8 as

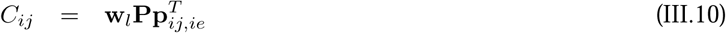

and the total cost for a given Dijkstra path 𝒫 as the sum of all individual edge costs for a given route:

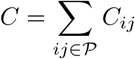

#### 4) Goodness of Fit

Given the cost function described in Equation III.8, we now need to develop a “goodness of fit” metric relative to the problem at hand. Referring back to Figure III.1, we are trying to find a weight vector, w_*l*_, that when used by Dijkstra’s algorithm with an edge cost defined by Equation III.8 will yield a path that is co-linear (or very-almost co-linear) with the original reference path specified.

Consider now Figure III.2 which shows part of a target path about which are successive colored regions. Each region is one vertex removed from its immediate inner region – hence they are referred *ISO*-regions. The target path occupies the inner-most region. About it, and at a distance of one vertex removed, is ISO Region 1. ISO Region 1 therefore defines a strip that is three vertices wide – the inner target path itself, and a single vertex wide strip on either side of this path. In a similar manner, the single-vertex region about ISO Region 1 defines ISO Region 2, and so forth until ISO Region 6, which is a ribbon patch 13 vertices wide with the original path at its very center.

**Figure III.2.**
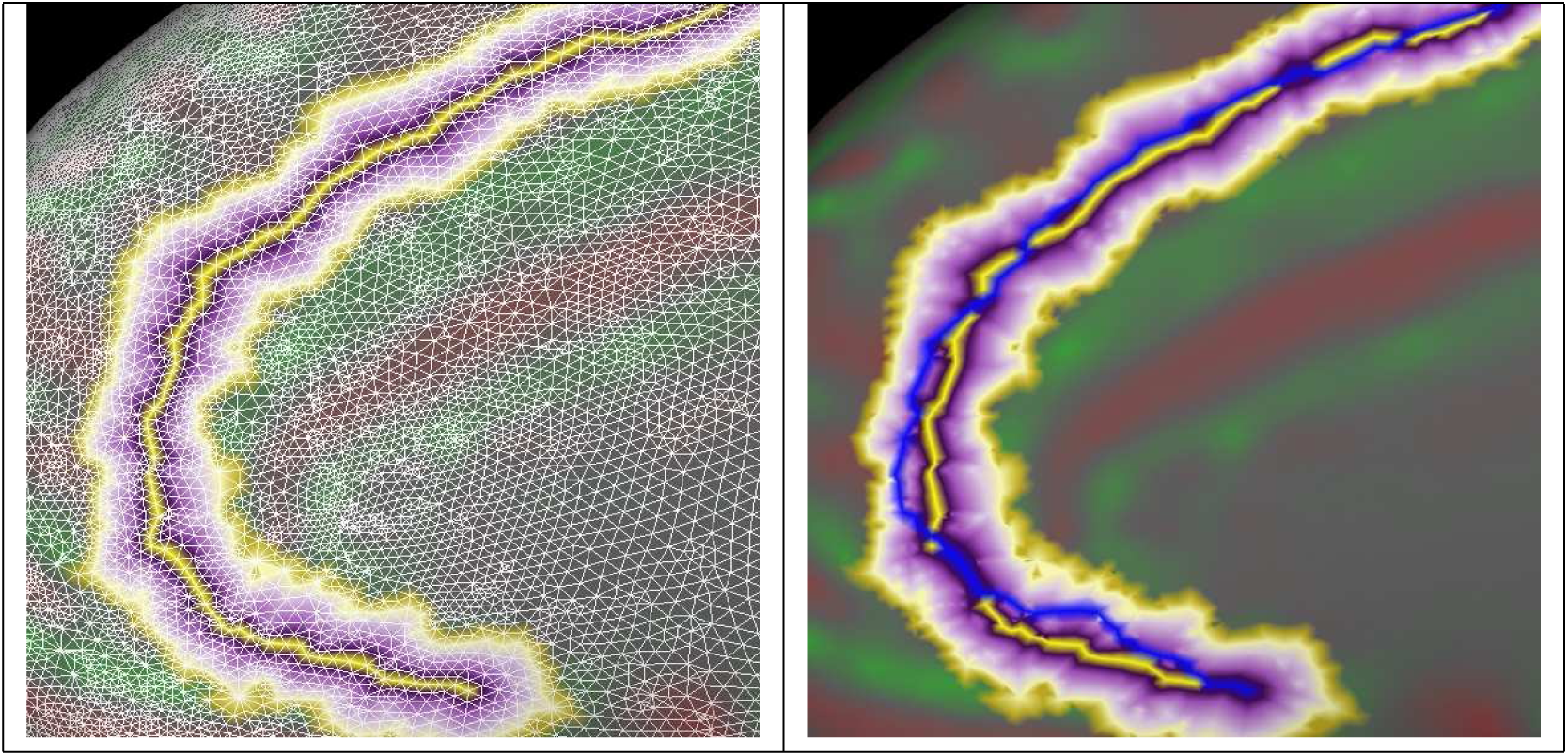
A series of ISO-regions about each target path were used to evaluate the utility of a candidate weight vector. On the left, the original path surrounded by its ISO Regions, shown projected on a wire-frame. On the right, a candidate path in blue that attempts to track the original path. Notice how different parts of the candidate path intersect different ISO Regions. The “fitness” of a candidate path is determined by how many vertex points along its length are outside successive ISO regions. Thus, an exact match would have 0 points outside all the ISO regions.

Conceptually, the most straightforward “goodness of fit” measure is simply the error distance between our reference signal and the Dijkstra generated signal. We can computationally simplify this measurement without needing to calculate distances, but consider instead an ISO region weighted function of the number of mesh vertices of the Dijkstra trajectory that lie in each ISO region. For a given ISO region *r*, we can express a percentage overlap of the Dijkstra path in this region as

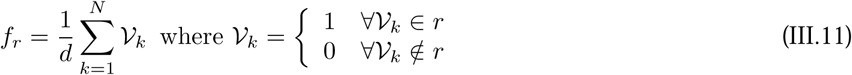

where *d* is the length of the original reference path, and 𝒱_*k*_ is a straightforward membership function of vertex *k* in the Dijkstra generated path for ISO region *r* about the reference path. Given this overlap fraction for ISO Region *r*, we can express the combined weighted overlap across all regions as

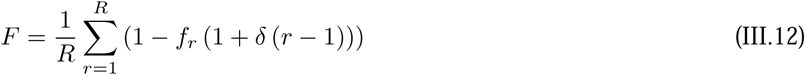

where *δ* is a weight penalty that penalizes overlap in higher ISO regions. This simpler calculation provides the same benefit to using a more complex distance metric: a Dijkstra path that meanders further away from the human reference has a lower overlap *F*. An example Dijkstra path is shown in blue on the right of Figure III.2, projected on the original path and with the ISO Regions shown. Notice how the Dijkstra path travels through several different ISO Regions.

### B. Weight vector optimization

The underlying nature of *C*_*ij*_ is highly non-linear in its greedy weight vector, w_*l*_ – in other words small *δ*w_*l*_ perturbations have no or little practical effect on *C*_*ij*_. This negated the use of gradient-based search methods. Moreover, the stated purpose of this paper is to describe sulci as a binary weight vector. This design decision considerably simplified our optimization regime – specifically for our nine-dimensional weight vector, a binary search needs only consider 2^9^ = 512 possible weight factors, which lay comfortably within the realm of exhaustive exploration. For each of the 512 possible weight vectors for a given sulcus on a given subject, we generated a shortest path using Dijkstra’s algorithm off the weight vector, and calculated the percentage overlap between the generated path and our reference using the weighted ISO method of Equation III.12. The weight vector with the highest overlap was chosen as the optimal for the particular subject and particular sulcus.

### C. Generalizing optimal weight vectors to templates

Each sulcus of each subject of each hemisphere can potentially have its own weight vector (or a set of weight vectors that map to the same “best” fitness value). One of the goals of this project was to determine if there is any “similarity” in terms of weight vectors for a given sulcus. In other words, is it possible to generalize features of a particular sulcus’s weight vector across all similar sulci from all the subjects?

One possible way to generate a template weight vector from a set of subject’s weights is to simply find the median value of all the relevant individual weight vectors, based on the assumption that this is the most representative sampling of the solution space. In order to evaluate the utility of this template with reference to the entire space of “best” vectors, we also perform a clustering based analysis to determine that the median vector lies within the larger “cloud” of solution vectors.

Such a clustering methodology is well established in statistical theory, and in fact is available as a set of tools and function calls in MatLAB. For this project, we considered each solution vector as a point in a nine dimensional space, and using a Hamming distance metric, generated a hierarchical cluster tree. We then partitioned the cluster tree using a cutoff value of 10, i.e. partitioned the space into a maximum of 10 clusters. The largest cluster was then determined, and a the overlap between the median template vector and this cluster was calculated, i.e. we determined how many bits were in common between the template and the largest cluster space of all the weight vector data. This overlap was used as a measure of confidence for the template vector’s validity.

## IV. Results and Discussion

For each of the twenty subjects in this study, ten sulci on each hemisphere were optimized. This resulted in 20 × 10 × 2 = 400 sulcal trajectories. We considered three conditions: (1) the trajectory fit for each sulcus as optimized to each subject; (2) the trajectory fit for each sulcus using a single template weight generalized from the optimal weights of each subject; and (3) the trajectory fit using a NULL (or zero) weight vector. If a given subject and given sulcus as parametrized by the weight vector exactly “fits” over the human generated example, the trajectory fit is 100%. An overview of all the results for each of theses conditions is show in Figure IV.1. The percentage overlap is shown per subject and per sulcus, with a blue-red colormap (bluer colors represent worse overlap, redder colors better overlap). Sulcal indices are as defined in the Figure.

**Figure IV.1.**
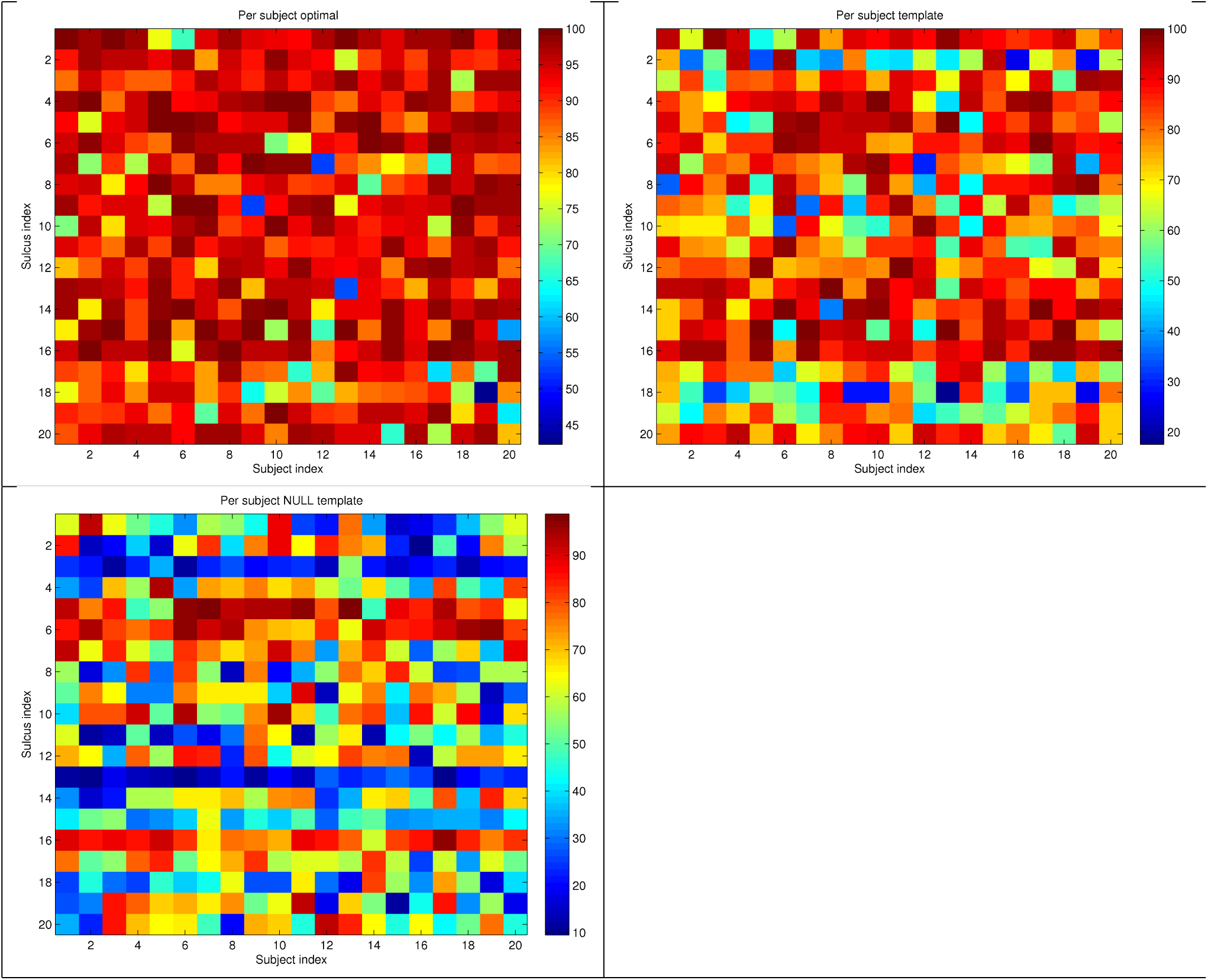
Percentage overlap of generated sulcal trajectories per subject and per sulcus. On top left, the overlap of the per subject, per sulcus optimals; on top right the overlap of the per sulcal group binary template weights; on bottom left the baseline NULL weight vector results. Subject indices are from left-to-right, and sulcal indices from top-to-bottom in order: 1: lh_calc; 2: lh_ce; 3: lh_ci; 4: lh_if; 5: lh_ol; 6: lh_phr; 7: lh_po; 8: lh_prc; 9: lh_sf; 10: lh_st; 11: rh_calc; 12: rh_ce; 13: rh_ci; 14: rh_if; 15: rh_ol; 16: rh_phr; 17: rh_po; 18: rh_prc; 19: rh_sf; 20: rh_st.

Figure IV.1 shows that the per-subject optimals are close to overlap with the human examples; the generalized sulcal templates had less good overlap; and the NULL weight baseline showed generally poor behavior. Note that we immediately see in the NULL template the poor performance of the cingulate (indices 3 and 13).

The results and discussion are organized into three sub-sections. First, we consider the system behavior on a per-sulcus manner evaluated across all patients. This is followed by a perspective taken from each subject considering all sulci for that subject. Finally, we present a blinded human expert evaluation of all sulcal traces.

### A. Overlap results from a sulcus perspective

Table II explores Figure IV.1 from the perspective of each sulcus. For each group (optimal, template, and NULL), three result columns are shown. In the “optimal” group, the weight values for a a single example subject and the percentage fit for this subject are presented, followed by the percentage fit averaged over all subjects. Thus, we see that for the “lh_calc” sulcus, one example subject had an optimal binary weight descriptor of “010000011” which resulted in a 99.68% weighted overlap. By comparison, taken across all subjects, the average for the optimal “lh_calc” sulcus weight resulted in a 94.20% overlap over the human reference (and each subject had a uniquely optimized sulcal weight). Also notable are some comparative observations evident between our chosen example and the whole subject group. For this example, the “lh_st” sulcus only optimized to a 70.50% overlap, and was the worst performing sulcus in this case. Taken across all subjects, however, this sulcus averaged 91.56% overlap. Clearly this example performed worse than the group average. The lowest overlap across all subjects was for the “rh_prc” at 81.48%, which also scored low for the subject shown. These observations might indicate that certain specific sulci optimize with more difficulty than others and might do so in a consistent manner.

**Table II.**
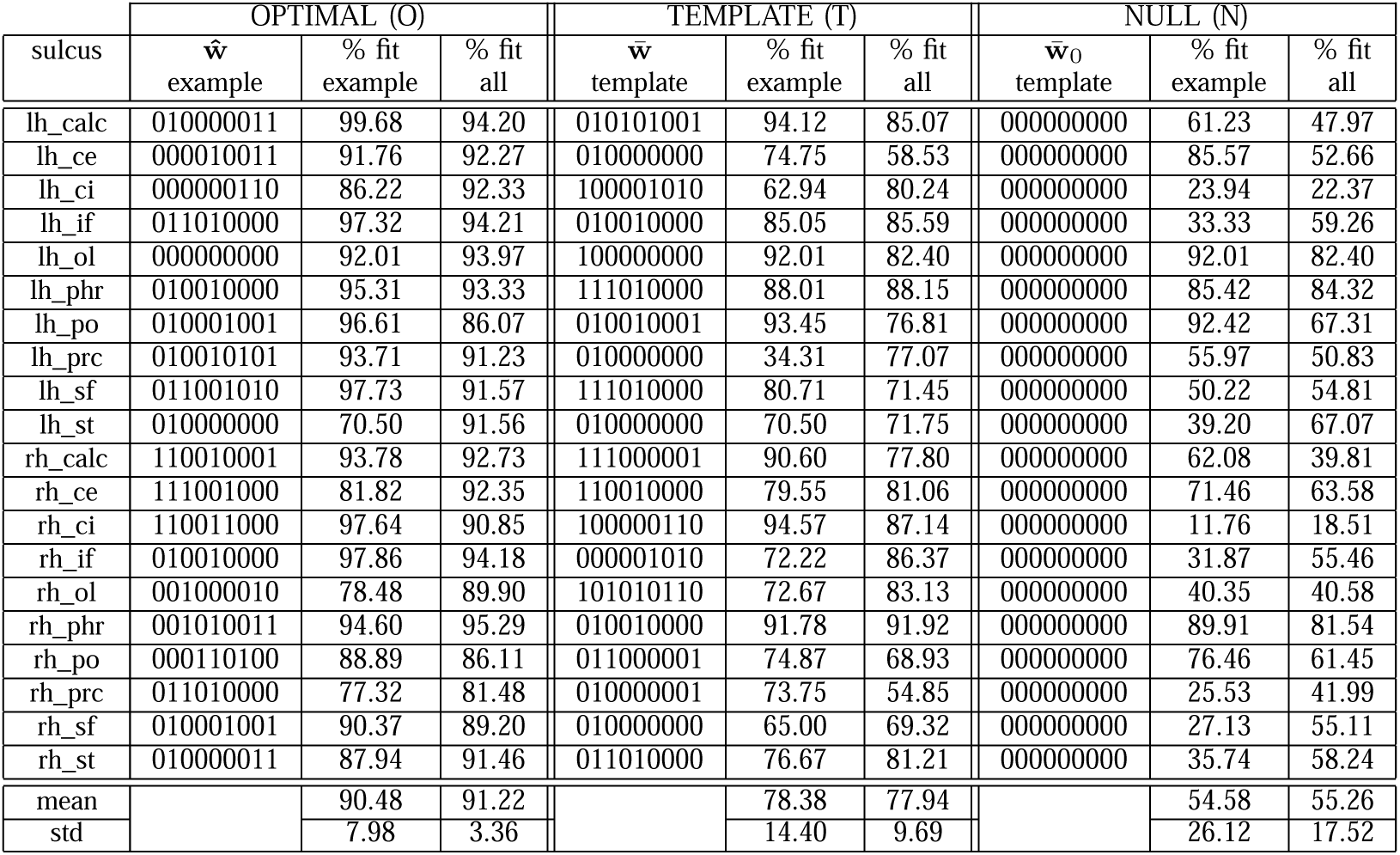
The optimal vs template vs null binary weight descriptor behavior for each sulcus. In the optimal case (columns on left), the binary sulcus descriptor for a specific sulcus is shown in ŵ followed by the percentage overlap this sulcus descriptor generated compared to the human reference in the “% fit example” column. The “% fit all” column shows the average overlap of each optimal sulcus for all the subjects (each subject had its own descriptor). In the Template columns (center), the template descriptor for each sulcus is shown in 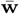. This same descriptor was used for each sulcus across all subjects. The “% fit example” shows the overlap that the template generated when used for an example case, followed by the average “% fit all” that this template resulted in across all subjects. In a similar fashion, the NULL Weight vector is shown on right for a specific example and across all subjects.

In the middle part of Table II, the template binary weight for each sulcus is shown, followed by the percentage sulcal trajectory overlap that this template resulted in for the same example subject as the “optimal” part of the table. The “% fit all” column shows the overlap of each template weight sulcus averaged over all subjects. Each sulcus now had the same template weight for all subjects, for example the “lh_calc” had a binary template of “010101001” which was applied to all “lh_calc” sulci on all subjects. Overall, the relative overlap percentages for the templated weights scored lower, which given that we have generalized away from a set of optimals matches our expectations. The deviation in overlap for the template weights, however, was lower than the subject-specific optimals, suggesting that generalization was preserving certain underlying properties of sulci across all subjects. Certain sulci, such as the “rh_prc” performed uniformly badly. The template binary resulted in an average 54.85% overlap across all subjects. It also scored comparatively low on the per-subject optimal shown in the first part of the table at 77.32% overlap, with also the lowest group optimal at 81.48%. Figures IV.2 provides a comparative “real world” references for the overlap percentages of Table II, with human traced sulcal references compared to the optimally weighted sulci on left, and also the human reference sulci compared with the templated binary sulci descriptions on right for the same subject.

**Figure IV.2.**
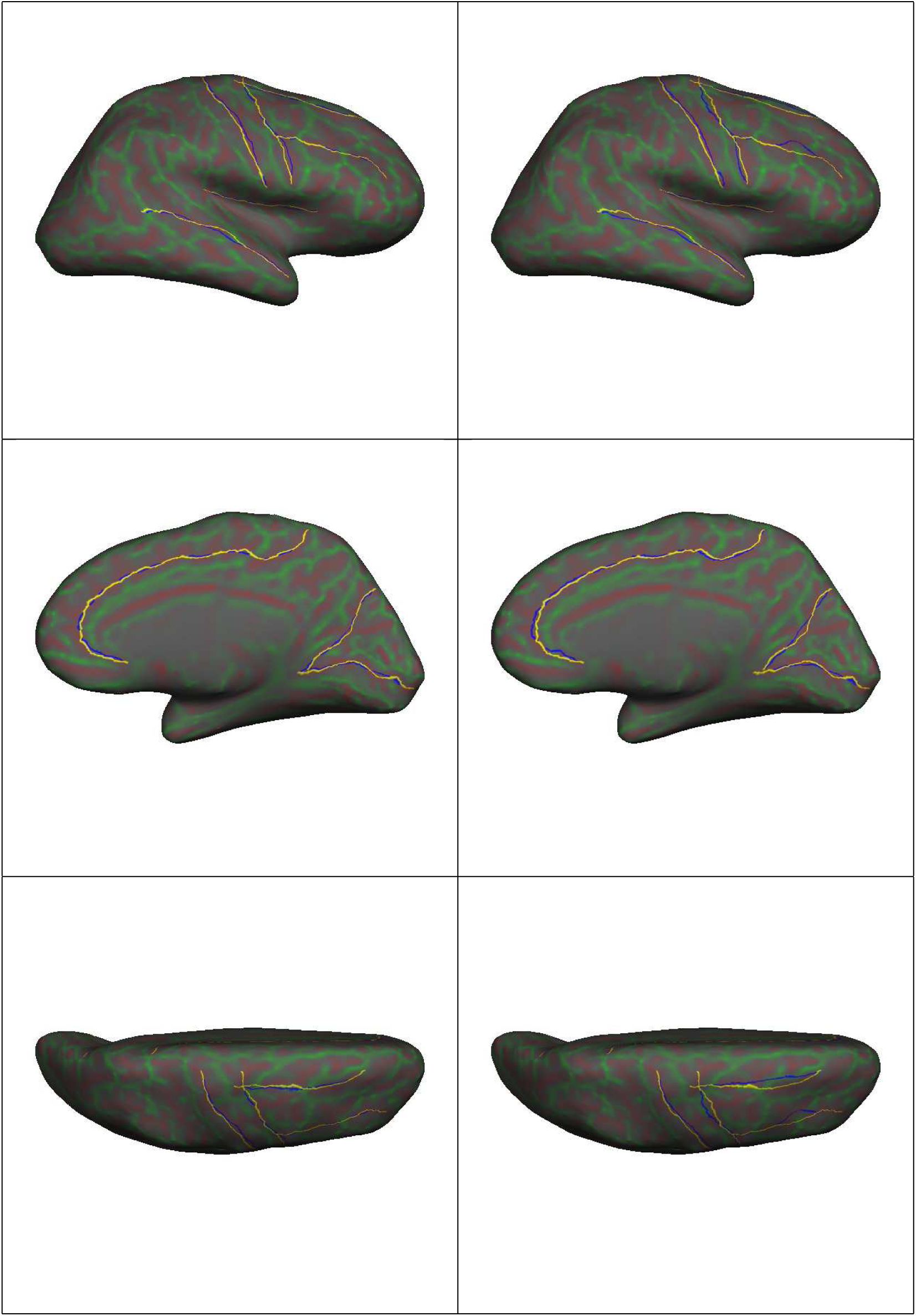
Human traced sulci (yellow) vs. binary weighted Dijkstra paths (blue) for the subject of Table II, and shown on the right hemisphere inflated surface. On left, lateral, medial, and superior aspects comparing the human references to the optimal binary descriptions. On right, human reference traces compared with the binary template description. The background coloration of red and green denote curvature values of the white matter surface projected onto the inflated reconstruction.

The final component of the Table shows the resultant behavior using the NULL template. In this case, all sulcus weights were set to zero, and the algorithm essentially showed no bias to any cost weight, and merely traveled along each mesh edge until it happened to connect the start and terminus vertices. The NULL templates are used to define a baseline reference against which the optimal and templated weights can be compared. The mean overlap in this case for all sulci across all subjects was 54.58%, with a 26.12% standard deviation. Note though how sensitive some sulci can be to weights. The NULL “lh_ce” sulcus had for the example shown in the table, a surprising 85.57% overlap with the human target. The template weight for this sulcus differed in only one bit, the *w*_*c*_ bit. This template weight for the example subject shown, dropped the overlap to 74.75% (and in fact, for this subject, the optimal “lh_ce” weighting was from a Hamming distance measure quite distal).

### B. Overlap results from a subject perspective

Using a similar organization as in Section IV-A, Table III presents overlap results organized per subject and shown across all sulci for that subject. Some additional information is also shown. Instead of weight vectors, we present the fractional degradation in performance: 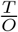 is the ratio of the Template overlap to the Optimal overlap, and 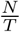 the ratio of the NULL overlap to the Template overlap. The product of these represents the overall degradation in overlap between the Optimal and the NULL weights.

**Table III.**
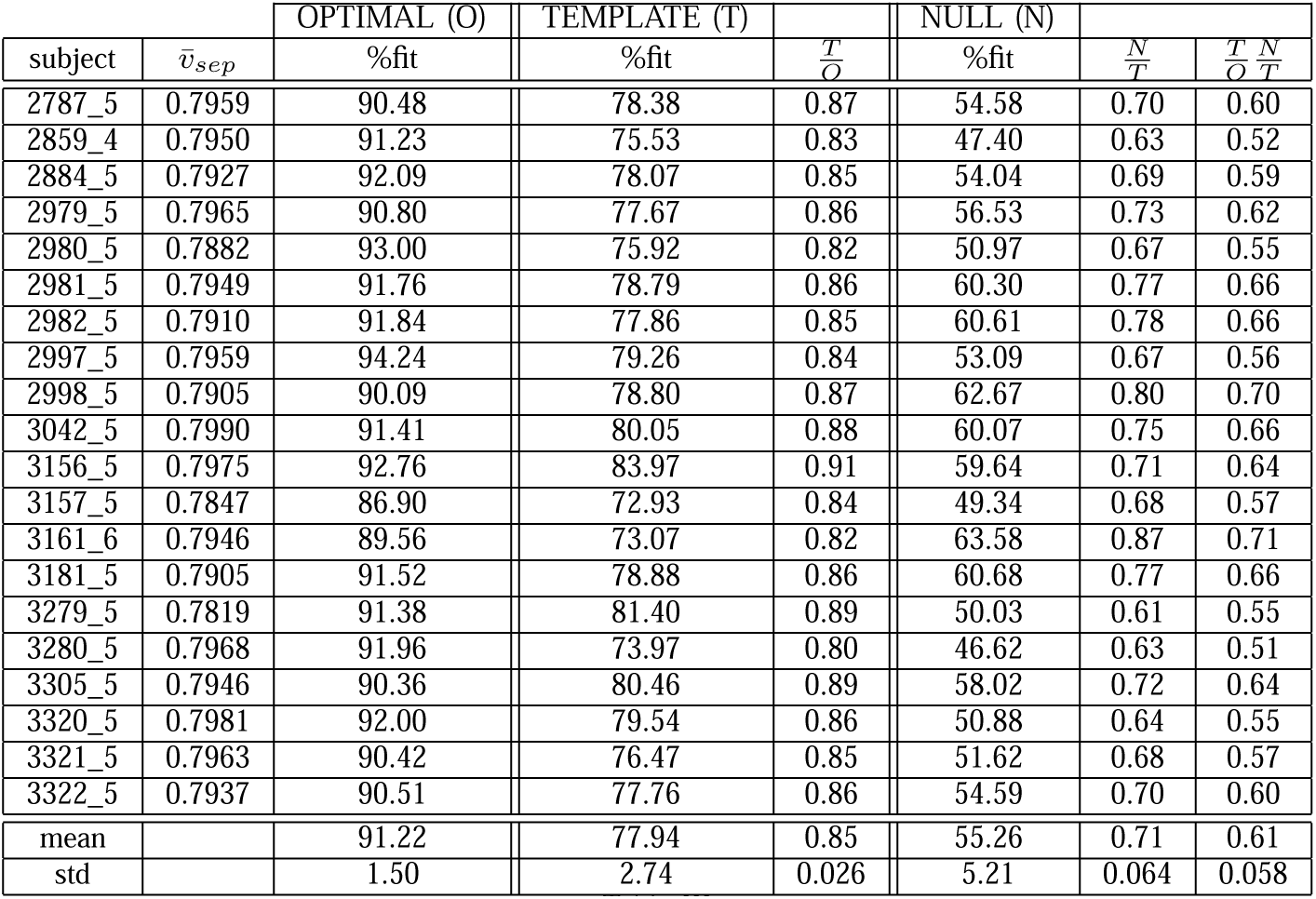
The optimal vs template vs null binary weight descriptor behavior for each subject. The 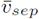 column indicates the average vertex separation for a particular subject. The optimal % fit column presents the results of all the optimal overlaps for a given subject averaged over all sulci for that subject. similarly for the template %fit and NULL %fit. The 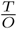 column shows the template fit as a percentage of the optimal fit. similarly for the 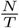 column. The final column presents the overlap of the NULL weight descriptor as a percentage of the optimal behavior.

As before, we note that the per-subject optimal weight vectors scored the highest overlap with the human reference sulci, with expected decreases when using sulcal template weights and a low scoring NULL reference. Taken over all subjects and sulci, the Template weights performed 85% as good as the optimal weights. The NULL weights performed 61% as well.

Subjects 2997_5 and 3156_5 scored the highest overlaps for their optimal weighting against their human generated references. In fact, subject 3156_5 scored the highest when using template weights, and (considering standard deviation) also high with the NULL reference weights – indicating that this specific subject’s sulci were perhaps geometrically more easily captured by this paper’s weight vector formulation and hence better conserved by the generalization process than the structural description of subject 3280_5 which had high optimal overlap, but comparatively low template and NULL weight overlap with its human traced sulcal references.

### C. Sulcal-template Optimizations

The template weights are an attempt to generalize the binary weight pattern for a given sulcus across all the subjects. Table IV shows summary results for each sulcus considered by this project. The first column presents the template weight vector, and the second indicates the correlation of this template to within the largest cluster of the space defined by all the individual weight vectors for that sulcus. In general we see that the correlation is 100%, except for 5 sulci that had 88% correlation, meaning that only 1 bit of the weight vector was not within the cluster space.

**Table IV.** Per-sulcus template weights, correlation, and deviation. The correlation column indicates what fraction of the weight template vector was contained in the largest cluster of points generated by all the optimal weight vectors for a particular sulcus. The remaining columns show the deviation for individual weight vector bits. The less deviation in a weight bit, the better the generalization of that bit.

| sulcus | $\bar{w}$ | correlation | $\sigma_P$ | $\sigma_d$ | $\sigma_c$ | $\sigma_h$ | $\sigma_{dc}$ | $\sigma_{dh}$ | $\sigma_{ch}$ | $\sigma_{dch}$ | $\sigma_p$ |
| --- | --- | --- | --- | --- | --- | --- | --- | --- | --- | --- | --- |
| lh_calc | 010101001 | 1.00 | 0.49 | 0.47 | 0.49 | 0.48 | 0.50 | 0.50 | 0.49 | 0.49 | 0.42 |
| lh_ce | 010000000 | 1.00 | 0.49 | 0.46 | 0.44 | 0.40 | 0.46 | 0.47 | 0.36 | 0.49 | 0.46 |
| lh_ci | 100001010 | 1.00 | 0.46 | 0.48 | 0.46 | 0.35 | 0.49 | 0.50 | 0.50 | 0.44 | 0.35 |
| lh_if | 010010000 | 1.00 | 0.47 | 0.34 | 0.36 | 0.00 | 0.50 | 0.49 | 0.00 | 0.50 | 0.44 |
| lh_ol | 100000000 | 1.00 | 0.50 | 0.45 | 0.42 | 0.46 | 0.49 | 0.47 | 0.40 | 0.50 | 0.49 |
| lh_phr | 111010000 | 1.00 | 0.50 | 0.49 | 0.50 | 0.39 | 0.50 | 0.31 | 0.43 | 0.40 | 0.48 |
| lh_po | 010010001 | 1.00 | 0.49 | 0.42 | 0.49 | 0.46 | 0.50 | 0.47 | 0.32 | 0.37 | 0.44 |
| lh_prc | 010000000 | 1.00 | 0.48 | 0.24 | 0.37 | 0.00 | 0.49 | 0.37 | 0.17 | 0.30 | 0.44 |
| lh_sf | 111010000 | 0.88 | 0.50 | 0.47 | 0.50 | 0.34 | 0.50 | 0.40 | 0.18 | 0.40 | 0.44 |
| lh_st | 010000000 | 1.00 | 0.50 | 0.47 | 0.44 | 0.50 | 0.49 | 0.49 | 0.30 | 0.22 | 0.48 |
| rh_calc | 111000001 | 0.88 | 0.48 | 0.46 | 0.50 | 0.48 | 0.47 | 0.29 | 0.46 | 0.32 | 0.46 |
| rh_ce | 110010000 | 1.00 | 0.49 | 0.48 | 0.49 | 0.38 | 0.48 | 0.50 | 0.35 | 0.50 | 0.50 |
| rh_ci | 100000110 | 0.88 | 0.43 | 0.39 | 0.45 | 0.36 | 0.43 | 0.50 | 0.45 | 0.48 | 0.00 |
| rh_if | 000001010 | 0.88 | 0.47 | 0.47 | 0.49 | 0.47 | 0.50 | 0.49 | 0.49 | 0.50 | 0.37 |
| rh_ol | 101010110 | 1.00 | 0.49 | 0.49 | 0.49 | 0.50 | 0.49 | 0.50 | 0.49 | 0.49 | 0.49 |
| rh_phr | 010010000 | 1.00 | 0.50 | 0.43 | 0.47 | 0.16 | 0.49 | 0.40 | 0.23 | 0.44 | 0.41 |
| rh_po | 011000001 | 0.88 | 0.49 | 0.49 | 0.49 | 0.45 | 0.49 | 0.17 | 0.45 | 0.28 | 0.49 |
| rh_prc | 010000001 | 1.00 | 0.44 | 0.37 | 0.43 | 0.35 | 0.49 | 0.15 | 0.15 | 0.15 | 0.50 |
| rh_sf | 010000000 | 1.00 | 0.49 | 0.50 | 0.41 | 0.40 | 0.47 | 0.45 | 0.47 | 0.46 | 0.35 |
| rh_st | 011010000 | 0.88 | 0.49 | 0.50 | 0.50 | 0.49 | 0.50 | 0.47 | 0.49 | 0.48 | 0.48 |
| mean |  |  | 0.48 | 0.44 | 0.46 | 0.37 | 0.49 | 0.42 | 0.36 | 0.41 | 0.42 |

It is important to note that the binary template description in and of itself is not a complete specification of a sulcus. Note in the table that the “lh_phr” and the “lh_sf” have the same template. While this does suggest that an underlying geometric similarity might exist between these sulci, each of these sulci are only fully specified in conjunction with their start and end vertices. Since Dijsktra’s algorithm is locally greedy, different start and end vertices for a sulcus, provided that they do not drift too much, would not change the overall sulcal weight description. We evaluated the effect of locally varying the start and end vertices on a sulcus’s weight optimization, and for variations within a range of 10 vertices, no changes in the optimal weights were observed.

The remaining columns of Table IV show the standard deviation of each particular weight bit from the median. For cases where the deviation is very close to 0.50, we can conclude that the underlying variation in that bit’s value ranged evenly between 0 and 1. Nonetheless, seen as a whole, some bits over the entire subject space showed less variation that others. In fact, the deviation in the *σ* _*d*_, *σ* _*h*_, *σ* _*dh*_, *σ* _*ch*_, *σ* _*dch*_, *σ* _*p*_ were all less than 0.45 while the remaining weights variations, i.e. *σ* _*P*_, *σ* _*c*_, *σ* _*dc*_ were all higher. We can interpret lower deviation values to imply a relative “bit strength” - the closer to zero deviation, the better the generalization of that particular bit.

### D. Human-expert evaluation

Although we can measure the deviation from the human-generated “gold standard” example templates of this “sulcus sink” system, a basic question remains as to how acceptable such generated sulcal paths are to a human expert. To this end, an independent expert anatomist who was not involved in the generation of the original sulcus paths, was presented with three sets of results for each hemisphere of each subject. Each set showed a complete grouping of projections of ten sulcal traces. One set showed the original traces as created by a human (the “target” set); a second set showed the subject-sulcus specific optimal, i.e. the best weight candidate for each sulcus on the particular subject as generated by the Dijkstra-based optimization (the “optimal” set); and a final set showed, for each sulcus, the trajectory resultant from a generalization of all weight vectors for each sulcus (the “template” set).

Without any knowledge of which set corresponded to which generation process, the evaluator was tasked with rating each sulcus of each set on a four-point (1 – 4) scale:

1. sulcus trajectory was completely unacceptable with significant portions of its path measurably far away from target
2. sulcus trajectory was unacceptable – typically the trajectory followed a neighboring, i.e. non-target, sulcus
3. sulcus trajectory was largely acceptable with portions of its path tracking along sulcal banks and not necessarily along the sulcal trough
4. sulcus trajectory was completely acceptable

Although there is of course a certain evaluator bias in such a rating system, generally speaking sulcal traces of “3” and “4” are acceptable, while those of “1” and “2” are not. Examples of these rating are shown in Figure IV.3.

**Figure IV.3.**
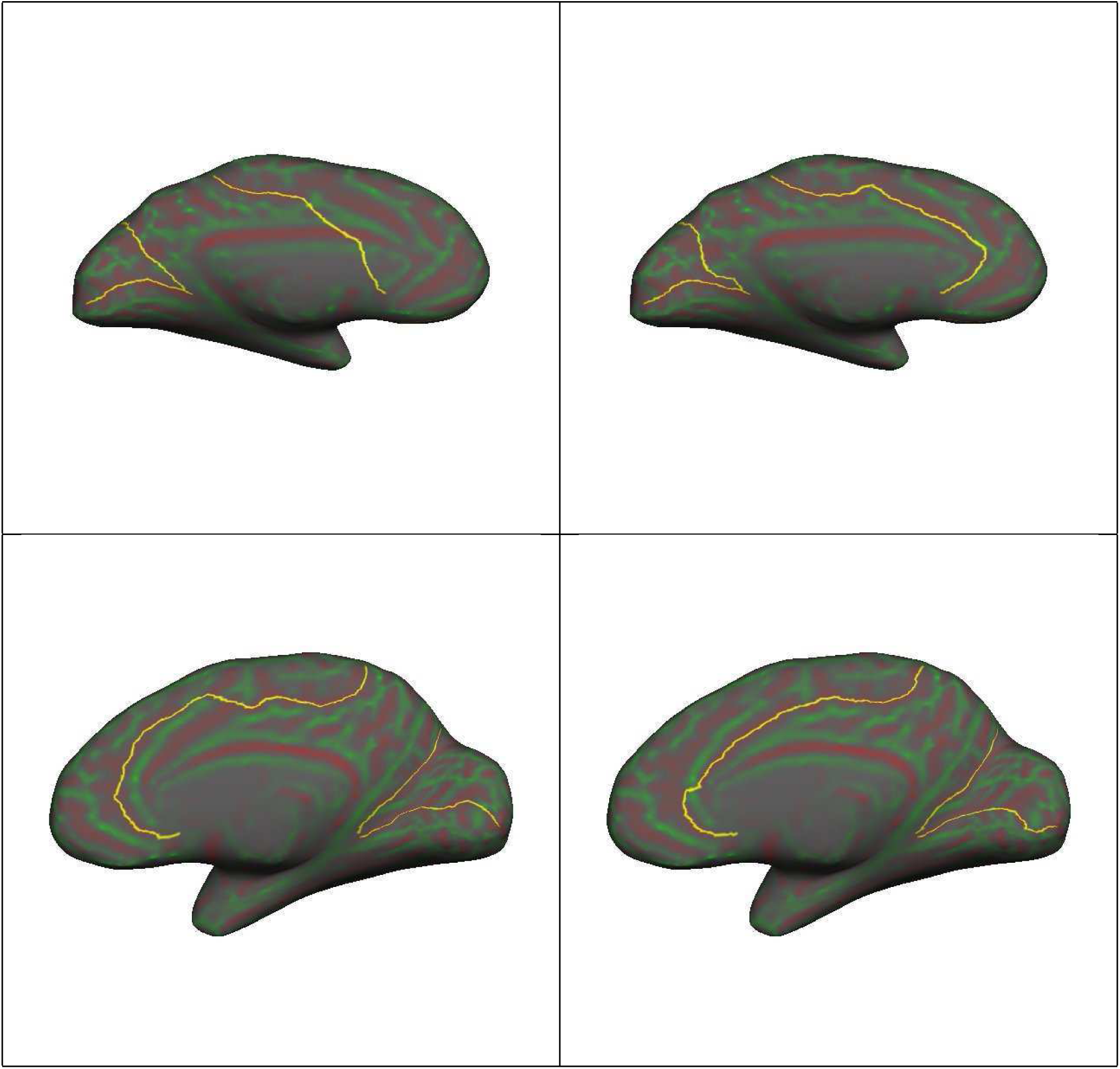
Examples of sulcal trace ratings using the cingulate sulcus (the long trace running length-wise along the medial aspect). Along the top row: “1” on left; “4” on right – notice that the “1” trace completely misses the anterior portion of the cingulate. Along the bottom row: “2” on left; “4” on right – in this case the “2” trace runs along the neighboring sulcus for most of its anterior component.

In the top row of Figure IV.3, the “1” rated cingulate was the result from using the generic cingulate template generated from all the optimal cingulate sulci of all subjects. The “4” rating on the right was the target trajectory as traced by a human. Interestingly, for the bottom row, the “2” rated cingulate on the left was in fact the trace as originally made by the human operator, and the “4” rated cingulate on the right was the same generic cingulate template that failed so completely for the top row subject.

Consider Table V which summarizes the sulcal trace evaluation on a per-subject basis. For each set of “target”, “optimal”, and “template”, the average rating for all sulci per subject per set was calculated. For the most part, the “target” set had the highest mean across all the subjects, followed by the “optimal”, with the “template” performing worst – a general trend that conforms with our expectations. Note however, some interesting deviations, particularly with the right hemisphere rankings. In some subjects (2859_4, 2884_5, 3156_5, and 3161_6) the trend for one or other hemisphere was reversed with the human target rated lowest, and the sulcus trace generated from the generic template rated highest. In these cases, several sulcal traces on these subjects made by the humans were not drawn well. How can the template perform better if some of its underlying domain set descriptors have errors? Examining these cases, we noted that only a small number of the sulci were traced badly. Our nine-bit binary descriptor had enough descriptive power to capture relevant features from the remaining larger pool of correctly drawn sulci to mitigate the errors in the training set.

**Table V.** Sulcus trace evaluations on a four point scale between 1 (worst) and 4 (best) as ranked by an independent human expert unaware of which sulcal set is the result of which process. Results are summarized on a per-subject basis, i.e. the average of all rankings for a subject/hemisphere/set.

| subject | Left Hemisphere |  |  | Right Hemisphere |  |  |
| --- | --- | --- | --- | --- | --- | --- |
|  | Human | Optimal | Template | Human | Optimal | Template |
| 2787_5 | 3.90 | 3.80 | 3.60 | 3.80 | 3.90 | 3.70 |
| 2859_4 | 4.00 | 3.90 | 3.90 | 3.60 | 3.60 | 3.80 |
| 2884_5 | 4.00 | 3.90 | 3.70 | 3.90 | 3.90 | 4.00 |
| 2979_5 | 3.90 | 3.70 | 3.70 | 4.00 | 4.00 | 4.00 |
| 2980_5 | 4.00 | 3.90 | 3.90 | 4.00 | 4.00 | 3.70 |
| 2981_5 | 4.00 | 4.00 | 4.00 | 3.80 | 3.80 | 3.80 |
| 2982_5 | 4.00 | 4.00 | 3.70 | 4.00 | 4.00 | 4.00 |
| 2997_5 | 4.00 | 4.00 | 4.00 | 4.00 | 4.00 | 3.90 |
| 2998_5 | 4.00 | 4.00 | 4.00 | 4.00 | 4.00 | 3.80 |
| 3042_5 | 3.90 | 4.00 | 3.70 | 4.00 | 3.90 | 4.00 |
| 3156_5 | 4.00 | 4.00 | 4.00 | 3.80 | 3.90 | 4.00 |
| 3157_5 | 3.90 | 4.00 | 4.00 | 3.80 | 4.00 | 4.00 |
| 3161_6 | 4.00 | 4.00 | 4.00 | 3.80 | 3.80 | 4.00 |
| 3181_5 | 3.80 | 3.80 | 3.80 | 4.00 | 4.00 | 4.00 |
| 3279_5 | 3.90 | 4.00 | 4.00 | 3.80 | 3.70 | 3.80 |
| 3280_5 | 3.80 | 3.80 | 3.70 | 4.00 | 3.90 | 3.80 |
| 3305_5 | 4.00 | 4.00 | 4.00 | 3.90 | 3.90 | 3.90 |
| 3320_5 | 3.80 | 3.50 | 3.50 | 4.00 | 4.00 | 4.00 |
| 3321_5 | 4.00 | 4.00 | 4.00 | 4.00 | 3.90 | 3.80 |
| 3322_5 | 4.00 | 4.00 | 4.00 | 3.90 | 3.90 | 3.70 |
| mean | 3.95 | 3.92 | 3.86 | 3.91 | 3.91 | 3.89 |
| std | 0.0759 | 0.135 | 0.167 | 0.115 | 0.110 | 0.118 |

Table VI organizes the summary rankings on a per-sulcus manner across all subjects. As with Table V, we note that the general trend is with the human “target” ranked highest, followed by the “optimal” track, with the “template” performing the worst. The human “target” also had the lowest score deviation, while the “template” sulcus had a measurably higher standard deviation.

**Table VI.** Sulcus trace evaluations on a four point scale between 1 (worst) and 4 (best) as ranked by an independent human expert unaware of which sulcal set is the result of which process. Results are summarized on a per-sulcus basis, i.e. for a given sulcus, the average across all subjects for a given hemisphere/set combination.

| sulcus | Left Hemisphere |  |  | Right Hemisphere |  |  |
| --- | --- | --- | --- | --- | --- | --- |
|  | Human | Optimal | Template | Human | Optimal | Template |
| calc | 3.900 | 3.90 | 3.91 | 3.90 | 3.90 | 3.85 |
| ce | 4.00 | 4.00 | 4.00 | 4.00 | 4.00 | 4.00 |
| ci | 3.90 | 3.75 | 3.05 | 3.90 | 3.95 | 3.65 |
| if | 4.00 | 3.85 | 3.91 | 3.90 | 3.90 | 3.90 |
| ol | 3.80 | 3.95 | 3.95 | 3.90 | 4.00 | 4.00 |
| phr | 4.00 | 4.00 | 4.00 | 4.00 | 3.95 | 4.00 |
| po | 3.90 | 3.90 | 3.91 | 3.85 | 3.85 | 3.85 |
| prc | 4.00 | 3.95 | 3.91 | 3.95 | 3.95 | 3.85 |
| sf | 3.95 | 3.90 | 3.95 | 3.95 | 3.95 | 3.95 |
| st | 4.00 | 3.95 | 3.95 | 3.70 | 3.60 | 3.80 |
| mean | 3.95 | 3.92 | 3.85 | 3.91 | 3.91 | 3.89 |
| std | 0.0685 | 0.0747 | 0.285 | 0.0864 | 0.117 | 0.11068 |

Note however, that in the “rh st” sulcus (right hemisphere superior temporal), and “rh/lh ol” (left and right hemisphere olfactory) cases, the human “target” was ranked lower than the “template” sulcus. In conjunction with Table V, this highlights subjects and sulci where the human operators might have made minor errors in creating the initial “target” traces for the system. For both hemispheres the “ci” (cingulate) sulcus had the lowest “template” rank – attesting to the fact that several “template” cingulate traces suffered from significant errors. This might suggest that the overall sulcus length might be an important factor in how well this system could generate optimal and hence template paths. The cingulate was by far the longest sulcus, and was consistently the lowest ranked.

## V. Conclusion

This paper described a methodology for compactly describing and tracing sulcal paths on reconstructed cortical surfaces. Using a sample set of 400 human-traced sulci paths, an optimization process attempted to find a weight vector that minimized the error between a sulcus target trajectory and a sulcus path generated by a Dijkstra-based search using the weight vector to define edge weights in the reconstructed cortical mesh. The system operated thus in a semi-automated supervised fashion since its optimization targets were provided *a priori* – and the optimization process itself was an *inverse* search since we were attempting to solve for weight factors that best mapped to a target path through Dijkstra’s algorithm.

The weight vector was a nine-element binary word that simply toggled on or off a set of contribution various features in the three dimensional surface space of a reconstructed brain. As such, the optimal weight vector for a specific sulcus represented along with the sulcus’s start and end points, a succinct binary parametrization or description of that sulcus.

Our approach with the methods of this paper is somewhat complimentary or tangential to the tendency to use highly automated methods. Fully automated methods tend to sacrifice a component of robustness while increasing repeatability and dramatically reducing analysis time compared to manual methods. To our knowledge, however, automatic sulcal extraction suffers from reliably identifying sulci and hence struggle to define starting and ending points of sulci. Moreover, automatic sulcal extraction methods do not only find major sulcal lines but also small and very minor sulcal lines (secondary or tertiary sulci) which can further complicate sulcal labeling and even introduce unwanted noise [21]. Sulcal lines can also be defined in wrong sulcal regions, or not defined in true sulcal regions [34, 35, 38].

During the training phase, optimal weight vectors per subject and per sulcus were found. Subsequently, a generalized binary “template” for each group of sulcal traces was created using a simple cluster median taken across all representative sulci. Thus, for example, all the right hemisphere cingulate weight vectors were analyzed to create a “template” right hemisphere cingulate weight vector. The underlying median and deviation for the group template bits were also analyzed for their relative stability – somewhat interestingly we found that the bit controlling curvature in the sulcus parametrization was less stable (with higher group deviation) than the bits weighing distance between neighboring vertices and sulcal depth. This suggests that sulcal descriptions might not be easily geometrically intuitive.

These template vectors represented thus a group-weighted description for an entire class of sulci (a full sulcus description was contained in the template binary weight and the start/end position of the given sulcus on a reconstructed surface mesh). The performance of these template vectors was measured, and compared with the per-subject optimals performed with an average of 80% overlap with about a 10% standard deviation (the optimal weights, relative to the human reference, had an average 91% overlap with a 3% deviation). We also considered so-called NULL templates with all weights set to zero as an absolute base reference. The NULL templates performed measurably poorly with only 55% average overlap with the human reference.

Finally, we presented a blinded human expert evaluator with sets of three sulci: one set contained the human target sulci for a subject, a second set contained the per-subject optimals as generated by this system, and a third set contained the sulcal paths generated using sulcal-template weights. Without knowledge of which set belonged to which process, the evaluated ranked individual sulci on a four point scale, with “1” being completely unacceptable and “4” completely acceptable. For the most part, the paths created by the template weights, though ranked consistently lower than the human targets or the optimal weights, were still quite acceptable to the evaluator.

Two interesting observations were made in the evaluator experiment. The first observation was that unacceptable paths were sometimes generated by the template weight for the cingulate sulcus, implying that a future automated system based on this work would still require special attention in the case of the cingulate. The second observation was that in a few cases, the human reference contained some errors as far as the evaluator was concerned. Interestingly, the general binary template sulcus weight, when applied to the same subjects containing human errors, were able to outperform the human reference. This indicates that enough information was generalized from the same sulcus that was correctly traced in other subjects.

Taken as a whole, the results presented here demonstrated that per-sulcus optimizations using an inverse Dijkstra-based method showed a very high degree of correlation between human target traces and automated ones. Moreover, generic template weight vectors generated from large sets of subject- and sulcus-hemisphere specific optimal weight vectors resulted, with the exception of the cingulate sulcus in a few cases, in trajectories that were acceptable to an independent human evaluator. We believe that the relative length of the cingulate sulcus compared to other sulci is a major reason for the poor performance in finding a binary description. The underlying geometry over such a long length could conceivably change in such a manner as to not be captured by our nine-element binary vector.

We have presented a method based on Dijkstra’s algorithm that can describe human sulcal paths with a nine-element binary weight vector. Per subject, per sulcus optimizations using this nine-bit binary description show on average a 91% overlap with reference human generated targets. Generalizations for groups of sulci based on per-subject optimizations perform well, indicating that such a binary description could be a useful and compact descriptor for sulci when coupled with a Dijkstra path optimizer.

Future work will seek to expand the weight vector with the addition of more parameters, including the Gaussian curvature and functions of the per-vertex principle curvatures. Since we were able to demonstrate good performance with a nine-bit binary description, it is our hope that performance can be measurably improved with straightforward (but still manageable) extensions to the search space size. In addition, more sophisticated generalization methods will be explored, and in the case of problematic cingulate sulcus, we propose to break this single sulcus into multiple joined segments, and optimize each segment independently.

